# Resistance to a CRISPR-based gene drive at an evolutionarily conserved site is revealed by mimicking genotype fixation

**DOI:** 10.1101/2021.07.26.453797

**Authors:** Silke Fuchs, William T. Garrood, Anna Beber, Andrew Hammond, Roberto Galizi, Matthew Gribble, Giulia Morselli, Tin-Yu J. Hui, Katie Willis, Nace Kranjc, Austin Burt, Tony Nolan, Andrea Crisanti

## Abstract

CRISPR-based homing gene drives can be designed to disrupt essential genes whilst biasing their own inheritance, leading to suppression of mosquito populations in the laboratory. This class of gene drives relies on CRISPR-Cas9 cleavage of a target sequence and copying (‘homing’) therein of the gene drive element from the homologous chromosome. However, target site mutations that are resistant to cleavage yet maintain the function of the essential gene are expected to be strongly selected for. Targeting functionally constrained regions where mutations are not easily tolerated should lower the probability of resistance. Evolutionary conservation at the sequence level is often a reliable indicator of functional constraint, though the actual level of underlying constraint between one conserved sequence and another can vary widely. Here we generated a novel gene drive in the malaria vector *Anopheles gambiae*, targeting an ultra-conserved target site in a haplosufficient essential gene (AGAP029113) required during mosquito development, which fulfils many of the criteria for the target of a population suppression gene drive. We then designed a selection regime to experimentally assess the likelihood of generation and subsequent selection of gene drive resistant mutations at its target site. We simulated, in a caged population, a scenario where the gene drive was approaching fixation, where selection for resistance is expected to be strongest. Continuous sampling of the target locus revealed that a single, restorative, in-frame nucleotide substitution was selected. Our findings show that ultra-conservation alone need not be predictive of a site that is refractory to target site resistance. Our strategy to evaluate resistance *in vivo* could help to validate candidate gene drive targets for their resilience to resistance and help to improve predictions of the invasion dynamics of gene drives in field populations.

**Author summary:** Gene drives have the potential to be applied as novel control strategy of disease-transmitting mosquitoes, by spreading genetic traits that suppress or modify the target population. Many gene drive elements work by recognising and cutting a specific target sequence in the mosquito genome and copying themselves into that target sequence allowing the gene drive to increase in frequency in the population.

Like other mosquito control interventions, efficacy will greatly depend on minimising the development of resistance to the gene drive mechanism - most likely via a change in the target sequence that prevents further cutting. One strategy to reduce resistance is to target sequences that are highly conserved, which implies that changes cannot easily be tolerated. We developed a strategy that simulates high selection pressure, under which resistance is most likely to emerge, and therefore provides a stringent test of its propensity to arise. Unlike previous results with another gene drive, we recovered a resistant allele within a few generations of gene drive exposure and at high frequency. Our results show that conserved sequences can vary hugely in ability to tolerate mutations and highlights the need to functionally validate future candidate gene drive target sites for their robustness to resistance.

## Introduction

CRISPR/Cas9 homing gene drives have shown much promise in providing an effective drive mechanism for novel genetic population modification and suppression strategies against insect vectors of major diseases, in particular the mosquito *Anopheles gambiae* (1–4). In general, these gene drives comprise a genetic construct containing a source of ubiquitously expressed guide RNA (gRNA) and Cas9 endonuclease that is expressed in the germline and together they cut a very specific chromosomal target site. Homology Directed Repair (HDR) leads to copying over of the entire gene drive construct into the target site in the homologous chromosome, leading to homozygosity in the germline of carrier individuals. Therefore, unlike Mendelian inheritance, potentially all offspring will receive a copy of the transgene, regardless of which chromosome they inherit from the parent. Modelling has suggested that suppression of disease vector populations could be achieved by designing the gene drive to target and disrupt haplosufficient genes (i.e. genes where one functioning copy is sufficient to maintain normal function) that are essential in the soma for mosquito viability or fertility, and confining the homing reaction to the germline (5–7). Under this scenario, individuals that contain one copy of the gene drive are ‘carriers’ that transmit the drive to a very high portion of the offspring, resulting in an increase in its frequency, which in turn places a strong genetic load on the population. This load is highest when all individuals carry at least 1 copy of the gene drive and the gene drive allele reaches genotype fixation,

Selection of mutations at the target site, or pre-existing alleles in the population, however, can hinder this type of gene drive strategy and prevent further spreading of the drive element (8). Mutations that prevent Cas9 cleavage at the intended cut site, whilst maintaining the function of the targeted gene, are referred to as ‘functionally resistant’ alleles. Such alleles can pre-exist in the target population or arise when the repair of the cleaved chromosome is mediated by non-homologous end joining (NHEJ) rather than HDR (8). One approach to reduce the development of resistance to a gene drive is to target regions that are structurally or functionally constrained and therefore less likely to tolerate insertions or deletions (‘indels’) and substitutions that could lead to resistance.

As a proof of principle, this approach has been used to mitigate resistance in small cage/scale testing observed in previous applications that targeted female fertility genes (2). By targeting a highly conserved 18bp sequence within the *A. gambiae doublesex* (*dsx*) gene, complete suppression of wild-type laboratory populations was achieved, while no alleles resistant to the gene drive were selected (1). Although these results are promising for the use of population suppression gene drives against these malaria vectors, it is important to have a range of suitable target genes, allowing a more tailored approach to achieve the desired levels of suppression and avoidance of resistance.

Our ability to identify genes that could be novel targets for gene drives has been greatly enhanced by genome resequencing efforts that have covered species across the *Anopheles* genus, spanning over 100 million years on an evolutionary timescale, as well as wild caught individuals of the closely related *Anopheles gambiae* and *Anopheles coluzzii* species (9–11). A recent analysis of genomic data from 21 *Anopheles* species revealed over 8000 ultra-conserved sites (100% identity) of sufficient length (18bp; corresponding to the minimal sequence recognised by a guide RNA) to be targetable by homing-based gene drives (12). It is to be expected that among this set of sequences there will be variation in their tolerance for changes to their underlying biological functions.

In this study, we focussed our analysis on ultra-conserved sites within putative haplosufficient essential genes as potential target sites for robust gene drives aimed at population suppression. Having generated an active gene drive targeting one of these essential genes, as proof of principle we subjected it to a maximal selection pressure to test its resilience to the selection of gene drive resistant alleles and its overall suitability for vector control.

## Results

### AGAP029113 is a haplosufficient essential gene in Anopheles gambiae

A recent analysis of ultra-conserved elements (UCEs) from 21 different *Anopheles* species and search of functional annotations of genes containing UCEs by O’Loughlin et al (12) identified ultra-conserved targets potentially affecting female fertility or lethal recessive phenotype that could be suitable targets for vector control. Amongst the list of potential targets, the *AGAP029113* gene is of interest as it contains multiple ultra-conserved sites with a total of 140 invariant sites. The *AGAP029113* gene is located on Chromosome 2L: 2895973-2931030 and was recently reassigned as a fusion of *AGAP004734* and *AGAP004732* (Vectorbase.org). It consists of 12 exons (Fig S1A) and contains domains putatively involved in interaction with the G-protein suppressor 2 pathway, DNA repair exonuclease subunit and DNA binding domains (Homeobox-like domain superfamily), suggesting a role in signalling, replication, recombination and DNA repair (13). Its high level of conservation and expression in larvae and in female ovaries (14,15), which does not change with increasing adult age (16) suggests that this gene is required during mosquito development in both sexes (15).

We chose a target site within AGAP029113 that is ultra-conserved across Culicidae (Culicidae; last common ancestor ∼ 150 million years ago) but not in *Drosophila melanogaster* genomes (Drosphilidae; 260 m.y.a.) (Fig. S1B). Even within *An. gambiae* and *Anopheles coluzzii* populations (2784 wild *Anopheles* mosquitoes sampled across Africa), this particular target site showed only negligible variations of less than 0.6% frequency (17) (Fig. S1C). This does not exclude the existence of further polymorphisms in wild populations but suggests that alteration of this sequence either naturally or by gene drive may have a strong fitness cost.

We then disrupted this candidate gene by inserting a GFP ‘docking’ cassette (‘*hdrGFP*’) into the target site within exon 5, which is expected to prevent the generation of a functional *AGAP029113* transcript and likely represents a null allele. The insertion of this cassette was performed by CRISPR-mediated HDR (Fig. 1A), using previously described methodology (1,2). After confirming the correct integration of the docking cassette (Fig. 1B) we crossed heterozygous individuals with each other and scored their progeny. Among larvae we observed heterozygous (29113^hdrGFP/+^) and homozygous (29113^hdrGFP/hdrGFP^) individuals at the expected Mendelian ratio (Fig 1C). However, individuals homozygous for the null allele failed to emerge to adulthood. During general maintenance of the strain we noticed no obvious fitness effects in individuals heterozygous for the null allele, suggesting that this gene may represent a suitable haplosufficient essential gene as a target for a population suppression gene drive strategy, whereby ‘carriers’ with a gene drive disrupting a single copy of the target gene need to be competitive to be able to pass on the gene drive to their offspring.

**Fig 1.**
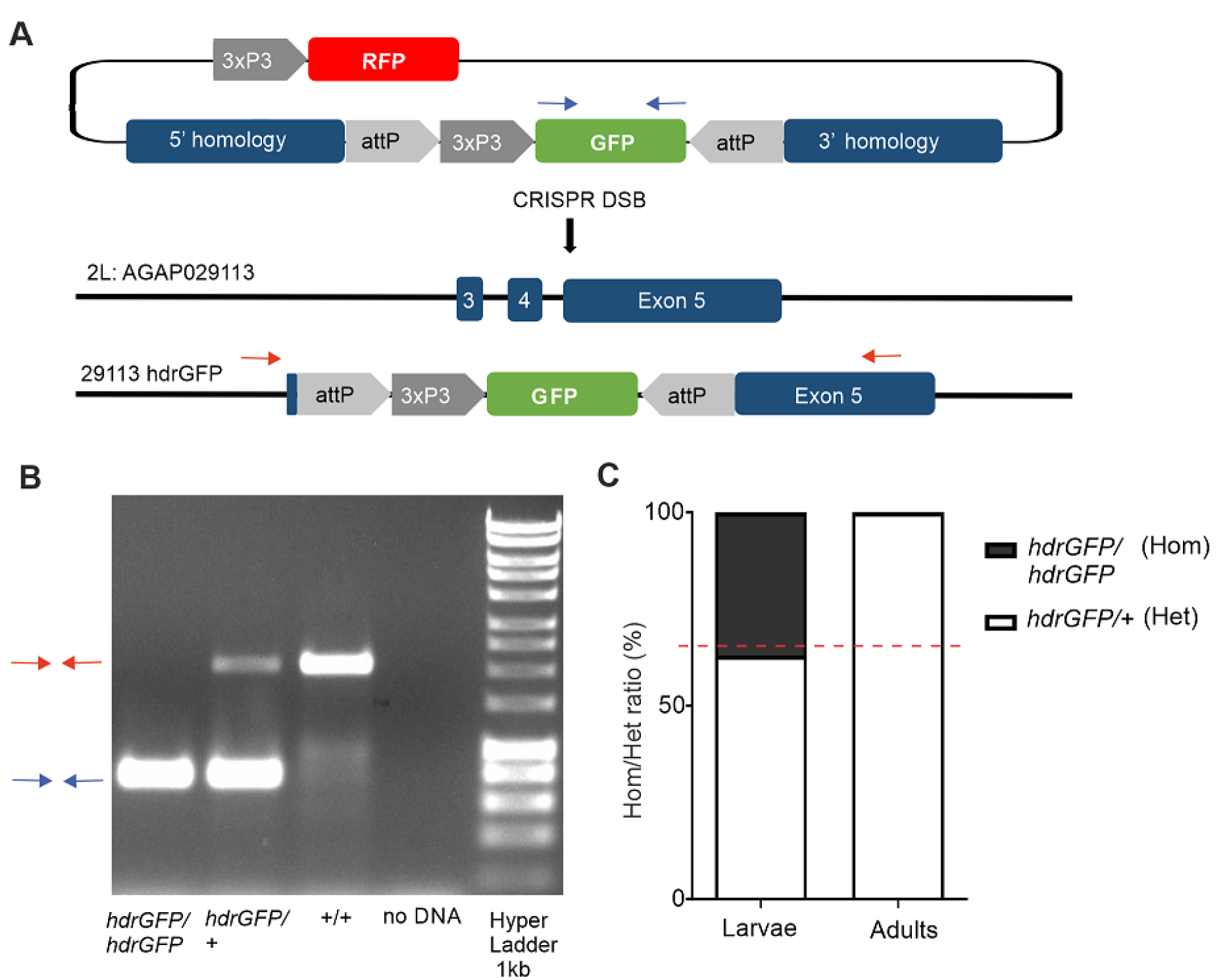
Gene disruption by HDR at a conserved site in exon 5 of *AGAP029113* causes recessive lethality. (**A)** Schematic representation of disruption of the *AGAP029113* gene. **(B)** PCR was used to confirm the targeted loci in WT individuals (red arrows) as well as those homozygous and heterozygous for the *29113*^*hdrGFP*^allele (blue arrow). **(C)** Heterozygous females and males were crossed with each other and the offspring was selected for GFP expression. Displayed is the percentage of homozygous (29113 ^hdrGFP/hdrGFP^) and heterozygous (29113 ^hdrGFP/+)^ L1 larvae (N=60) and adults (N=60) analysed by PCR as well as the expected homozygote vs heterozygote ratio according to Mendelian inheritance (red dashed line).

### Phenotypic assessment of a gene drive targeting an essential gene

To assess the phenotype in the presence of a gene drive construct targeting this locus, an active Cas9/gRNA cassette (29113^CRISPRh^) was inserted into the locus as described previously (5). The 29113^CRISPRh^ construct contained *Cas9* with expression controlled by the *zero population growth* (*zpg*) germline promoter (1), together with a dominant *RFP* marker gene to assist in tracking the inheritance of the gene drive (Fig. 2A). This was placed in locus via recombinase mediated cassette exchange (RMCE), replacing the GFP cassette with the 29113^CRISPRh^ cassette. This gene drive construct showed high rates of biased inheritance from heterozygous (29113^CRISPRh^/+) female (92.7% ± 2.9) and male mosquitoes (91.3% ± 3.2) (Fig. 2B).

**Fig 2.**
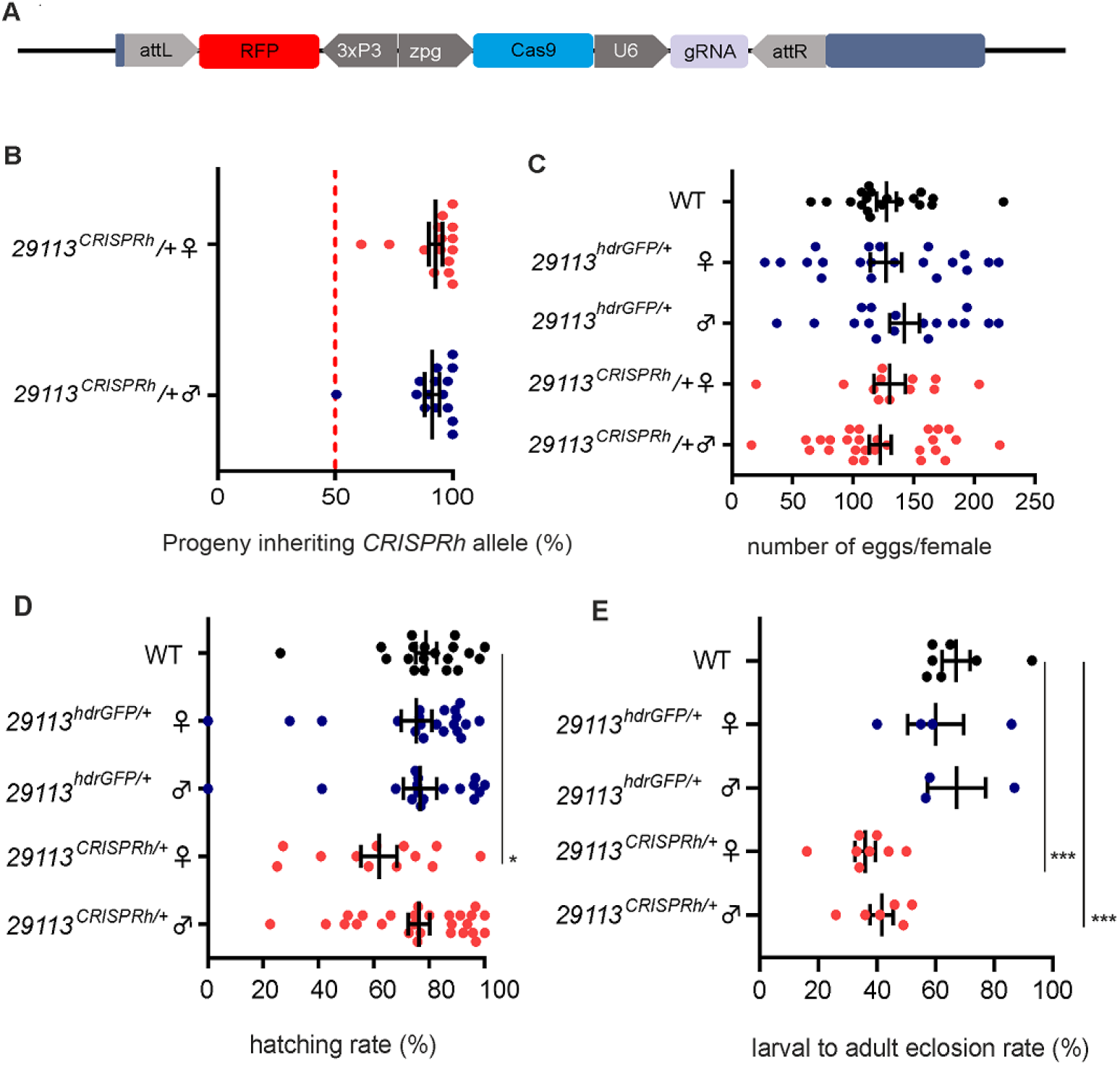
Phenotypic assessment of 29113^CRISPRh/+^ individuals crossed with wild-type. **(A)** Displayed is the CRISPR homing construct 29113^CRISPRh^ consisting of a *3xP3::RFP* marker, *Cas9* under the transcriptional control of the *zpg* promoter and a gRNA under the control of the ubiquitous *U6* PolII which replaced the GFP transcription unit in heterozygous 29113^hdrGFP/+^ lines by RCME. The gRNA cleaves at the homologous wild-type allele at the target site corresponding to the 29113^CRISPRh^ insertion. Repair of the cleaved chromosome through HDR leads to copying of the *CRISPR*^*h*^ allele and homing. **(B)** Male or female 29113 ^CRISPRh^/+ heterozygotes were mated to wild-type. High levels of homing of >90% was observed. Progeny from individual females were scored for the presence of the RFP linked to the 29113^CRISPRh^ construct and the average transmission rate indicated by vertical bar. A minimum of 15 females were analyzed for each cross. The average homing rate ± s.e.m. is also shown. **(C)** Scatter plots displaying number of eggs laid by single females in crosses of 29113^*CRISPRh*^*/+* mosquitoes with wild-type individuals. Data were analyzed by quasi-Poisson GLM model. No significant differences were found between the wild-type and transgenic strains. **(D)** The hatching rate of these crosses is also displayed. Data were analyzed by random effect GLM model with binomial response. No significant differences were found. **(E)** Larval to adult eclosion rate. Data were analyzed by random effect binomial GLM model. * p<0.05, ** p<0.01, *** p<0.001.

Impaired fecundity and fertility of heterozygotes has been observed in other gene drive strains (1,2). We performed fecundity assays where heterozygous (29113^CRISPRh^/+) males and heterozygous (29113^CRISPRh^/+) females were crossed to wild-type G3 mosquitoes and the number of eggs laid and hatched larvae was counted by individual oviposition of females. The wild-type G3 strain was used as control. There was no significant difference in the number of eggs across all 5 crosses (Fig. 2C; quasi-Poisson GLM, F=0.458, df1=4, df2=90, p=0.766). However, the hatching rate of the progeny of 29113^CRISPRh/+^ females was reduced by 21.6% relative to wild-type controls (Fig. 2D; z-test, p=0.046). There was also increased mortality during larvae to adult development of about 37.8% and 46.2% for heterozygous male and female offspring respectively (Fig 2E, z-test, 29113^CRISPRh^/+ males p = 3.33e-5; 29113^CRISPRh^/+ females p = 4.88e-8).

### Assessment of 29113^CRISPRh/+^ spread in a cage population experiment

Despite the fitness costs apparent with this gene drive in heterozygous carrier individuals, the extent of biased inheritance was still expected to cause the element to increase in frequency in a population (1,2). To test this, we seeded 29113^*CRISPRh*^/+ individuals into a wild-type caged population (total initial population size of 600) at a starting frequency of 20%, in triplicate, and monitored the frequency of gene drive individuals over time. The same experiment was performed in parallel with the non-driving 29113 ^hdrGFP/+^ mosquitoes as control.

For releases with the control strain the frequency of individuals with the control allele decreased slowly over 3 generations to an average of 10.6%, consistent with the expectation for a non-driving null allele (Fig. 3A). However, in the gene drive release cages, despite observing an initial increase in the frequency of gene drive carriers in the generation after release, this rapidly declined, to the extent that by 3 generations post-release the gene drive was lost from the population in all three replicates (Fig. 3B). Rapid loss of the gene drive suggests that there is a fitness cost associated with the Cas9/gRNA expression from the gene drive allele that prevents its persistence. We therefore used a simple deterministic model of a single, randomly mating population with two life stages (juveniles and adults), two sexes and discrete non-overlapping generations, following the structure of Beaghton et al (18) and incorporated the heterozygote fitness costs identified from the phenotypic assays. They included a) no fitness costs (grey dashed line) b) fitness costs that were observed in the phenotypic assays (21% reduced hatching rate in 29113^CRISPRh/+^ females and reduced larvae to adult eclosion rate of 37.8% and 46.2% for 29113^CRISPRh/+^ males and females) (light red dashed line) and c) further potential fecundity and somatic costs (90% for 29113^CRISPRh/+^ males and females) that were not measured (dark red dashed line). Overall, the data fitted the model c) assuming additional fitness costs associated with the gene drive construct. This could include fitness costs that manifest in adulthood which we did not assess.

**Fig 3.**
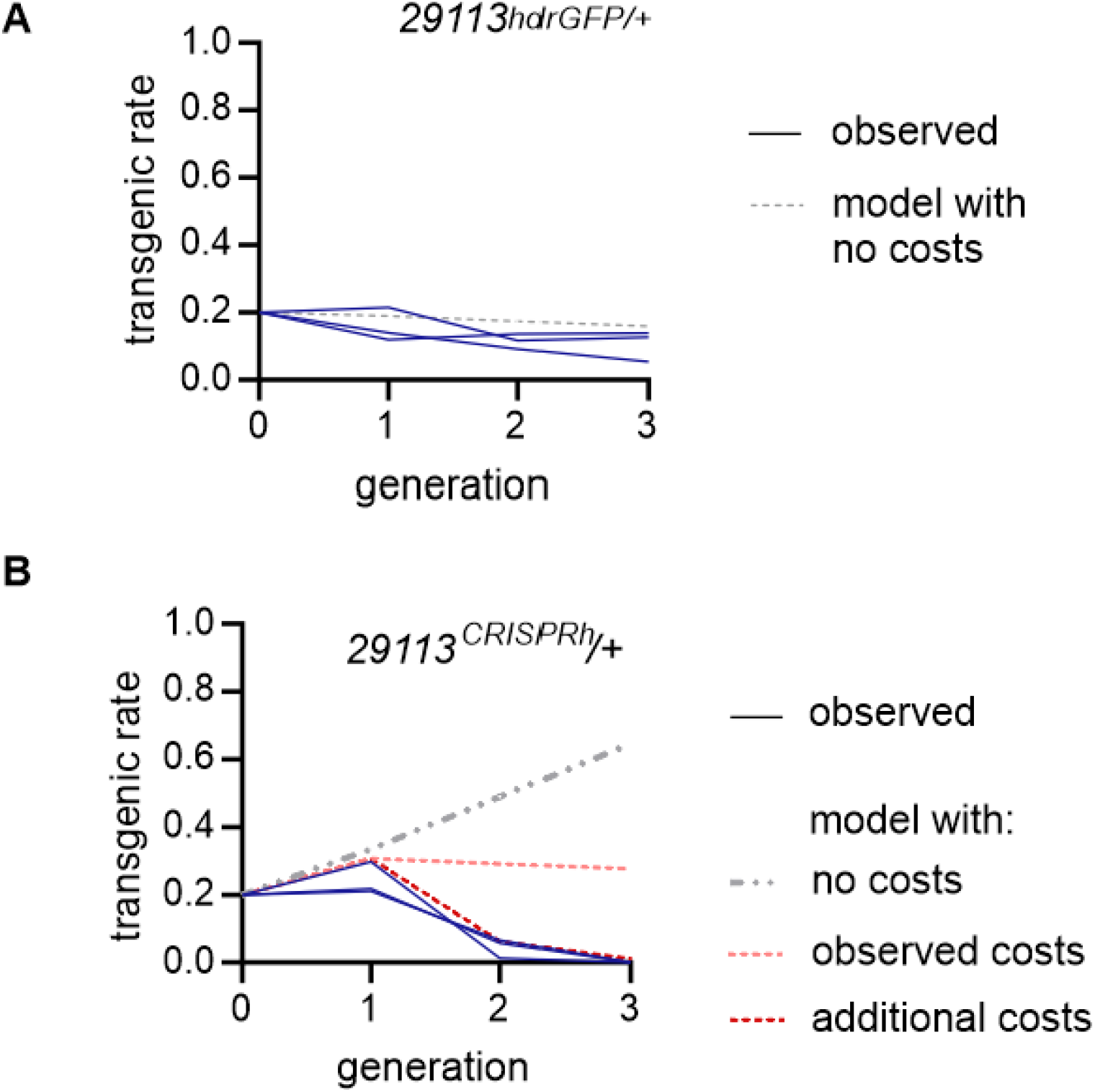
Frequency of individuals containing the 29113^CRISPRh^ drive allele over time in a population. Transgenics were introduced at a 20% frequency in a wild-type (G3) population. The transgenic rate was recorded in each subsequent generation. The experiments were performed in triplicates. **(A)** Proportion of individuals carrying the non-homing 29113^hdrGFP^ transgene (in blue). The dotted grey line shows the expected transgenic rate, based on no fitness costs in heterozygotes **(B)** Proportion of the 29113^CRISPRh/+^ individuals (in blue). Dotted lines show the deterministic prediction based on the observed homing rates for 29113^CRISPRh/+^ individuals assuming either no fitness costs in gene drive carriers (grey hatched line), observed fitness costs (light red line) from the phenotypic assays (fecundity cost (1 - relative hatching rate) (F) = 0.216 and somatic cost (1 – relative eclosion rate) (M) = 0.378, (F) = 0.462,) and additional somatic costs (M) = 0.9, (F) =0.9, in dark red line) not measured in our experiments.

We also investigated the possibility of the generation and subsequent selection of a functional resistant allele to the gene drive that could have led to loss of the transgene. Such alleles can be formed when Cas9-mediated cuts are repaired by error-prone end joining DNA repair pathways producing small insertions or deletions or substitutions (8). Our models showed that such resistant alleles would have had no impact on the transgenic rate over the first 5 generations of the population experiment (Supplementary Fig. 2). This suggests that the dominant factor in determining the drive loss was the unexpected additional fitness costs associated with harbouring the drive element in heterozygosity, rather than the emergence and selection of resistance. Nonetheless, an important question is the resilience to resistance at this target site, in the face of a gene drive.

### Non-drive alleles generated at the target site by Cas9 nuclease activity

To investigate the array of target site mutations that can be generated at the ultra-conserved site we devised a cross that enriched for non-drive alleles exposed to the nuclease from the gene drive (Fig 4A). In this assay the non-drive alleles are balanced against a marked (GFP+) null allele of the haplosufficient target gene. Consistent with an essential role for the *AGAP029113* gene between larval and adult transition, the distribution of in-frame and out-of-frame alleles was markedly different between the larval stage (where a complete null genotype is tolerated) and the adult stage (where it is not): pooled sequencing of the target site in L1 instar larvae revealed that 26% of non-drive alleles contained a target site mutation, of which 10.8% were in-frame (Fig 4B); sampling at the adult stage, only 5.7% of non-drive alleles contained mutations, of which 99.3% were in-frame. The persistence of in-frame alleles in the adult stage strongly suggests that these restore function, at least partially, to the essential target gene. If these mutations are also resistant to further cleavage by the gene drive then they would meet the criteria of being functionally resistant and thus be positively selected in the presence of the gene drive.

**Fig 4.**
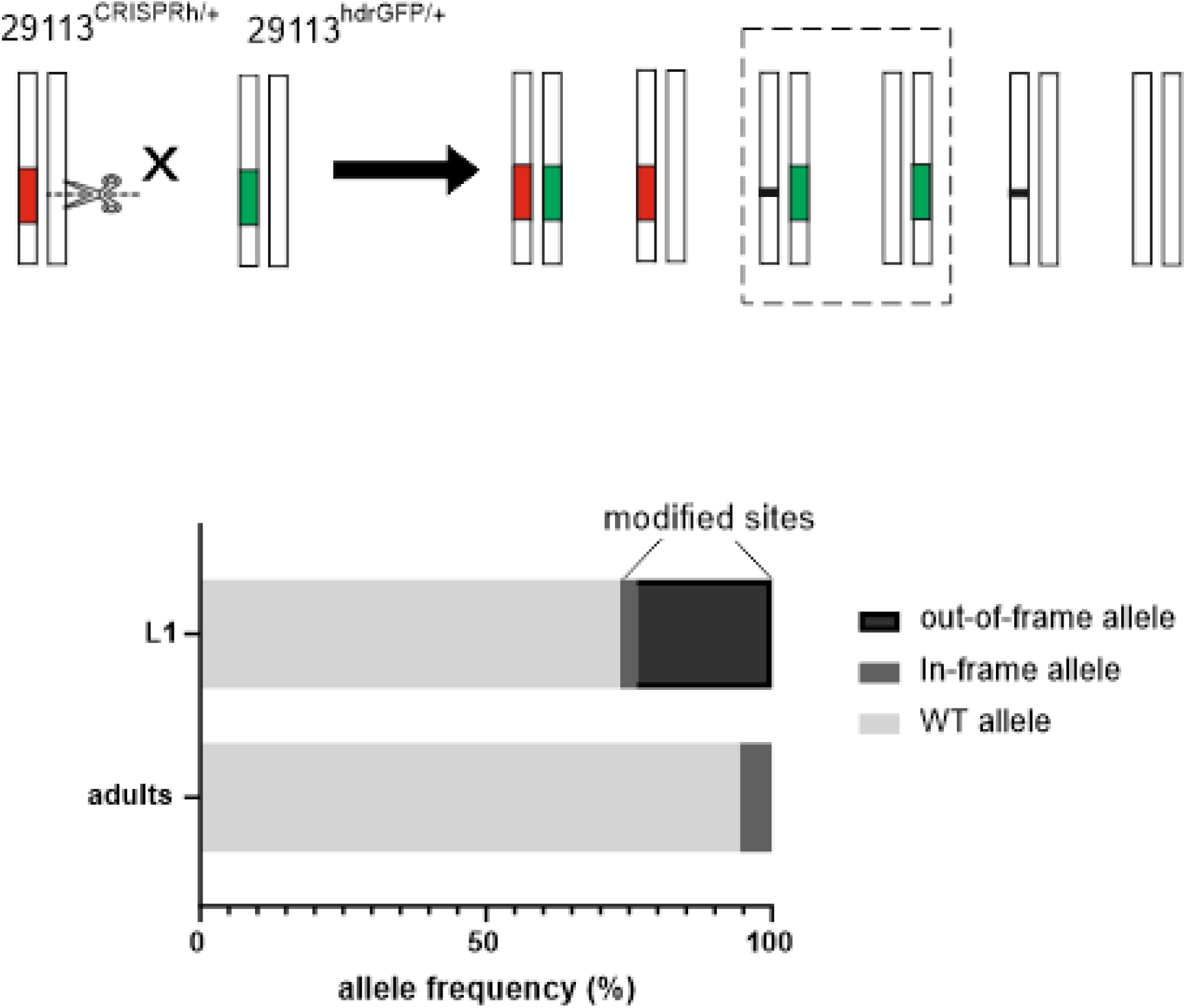
Resistance analysis of an ultra-conserved target site in *AGAP0029113*. **(A)** Crossing scheme devised to allow analysis of alleles that were exposed to CRISPR/Cas9, but which were not converted to the 29113^CRISPRh^ construct. The crossing of heterozygous 29113^CRISPRh/+^ individuals to 29113^hdrGFP/+^ individuals, and subsequent screening of the offspring for GFP+, provided mosquitoes that had an 29113^hdrGFP/+^ genotype, with the other allele being wild-type or containing indels generated by NHEJ repair. Red represents the 29113^CRISPRh^ allele (RFP+), whilst green shows the 29113^hdrGFP^ allele (GFP+) and black shows novel mutations. Progeny was screened for GFP+/RFP-(dashed rectangle). **(B)** Allele frequency (%) at the CRISPR/Cas9 cleavage site. L1: 7000 larvae, adults: 90 individuals.

### Selection of 29113^CRISPRh^ resistant alleles by mimicking genotype fixation in a caged population

In order to investigate our assumption that these mutations could be strongly selected, we mimicked a scenario of genotype fixation (i.e., 100% of mosquitoes have at least one copy of the gene drive), where selection pressure for functionally resistant alleles is expected to be highest. We crossed individuals heterozygous for the gene drive construct, allowing all offspring to survive and reproduce for multiple generations. Given our previous estimate of gene drive transmission rate (93%), under this scenario the vast majority (87% (0.93^2^)) of progeny would be expected to be homozygous for the gene drive and so would die during larval development. Of the progeny that survive to form the following generation, the majority would be heterozygous and destined to produce predominantly non-viable offspring. Under these conditions a functionally resistant allele will therefore have a high selective advantage since it will confer viability on any genotype that inherits it.

Over the first five generations of this experiment, the gene drive population (marked in red) was constantly suppressed (Fig. 5A), producing very few eggs compared to the non-drive 29113^hdrGFP/+^ controls (in green), consistent with the majority of the offspring lacking a functional copy of the *AGAP029113* gene. However, by generations 6 and 7 we observed a rise in frequency of larvae as well as the number of adults and eggs produced by the gene drive cage. Together, these observations are consistent with a breakthrough of functionally resistant alleles that are refractory to gene drive invasion and restore the carrying capacity of the population. Indeed, sequencing of adults from the G7 generation (Fig. 5B) showed over 87% of reads contained the same mutation at the Cas9 target site. We also detected this single nucleotide substitution amongst other mutations recovered in larvae and adults in our previous screen (Supplementary Fig. 3). Since this mutation was not found in the laboratory wild-type population (despite sequencing of 360 wild-type individuals and alignment of 35817 reads), the most likely explanation is that it is a de novo mutation caused by end-joining repair following nuclease activity of the gene drive. This specific mutation was selected and rapidly increased in frequency over generations likely due to a fitness advantage compared to other indels found in the single generation assay.

**Fig 5.**
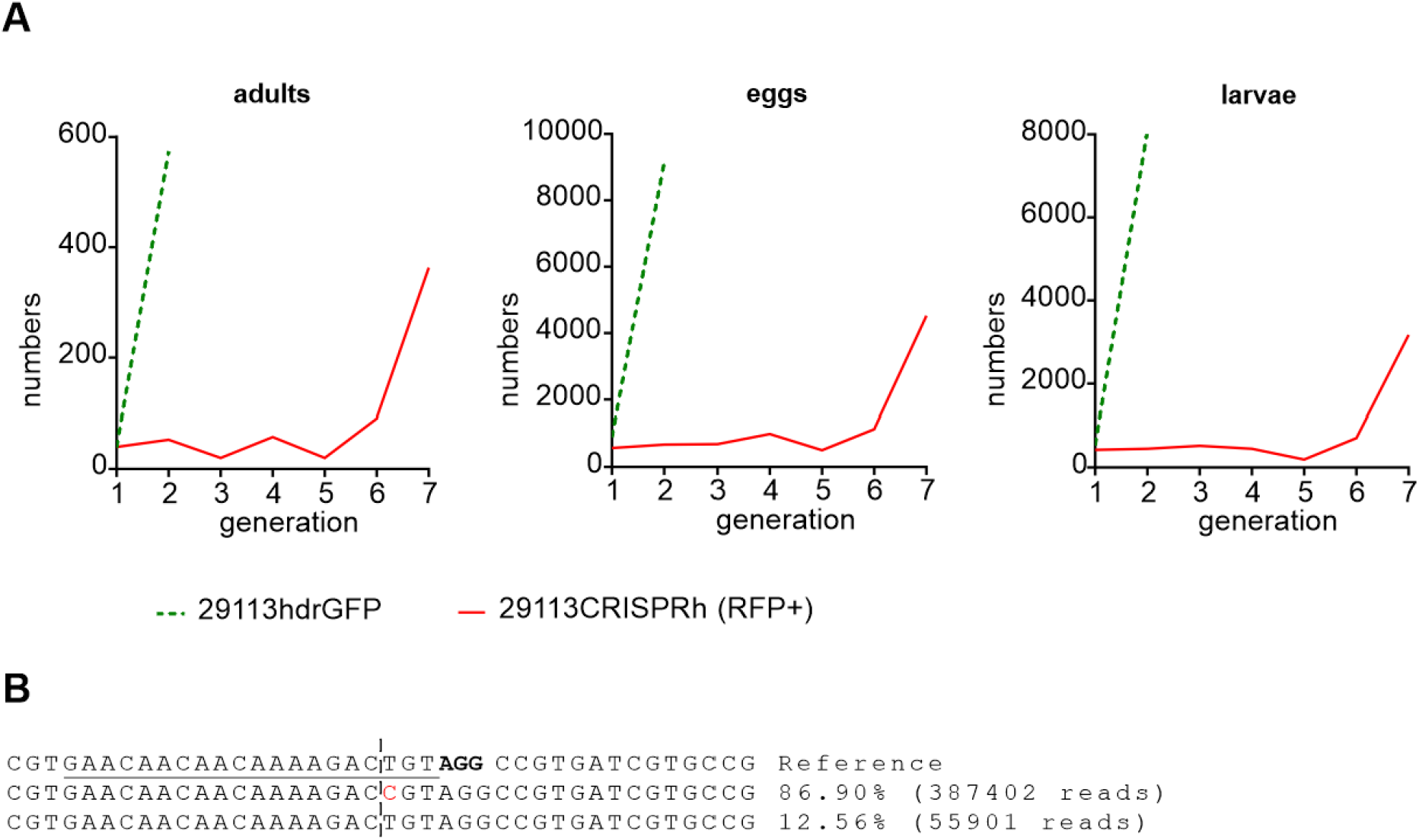
Reproductive output of a population when mimicking near genotype fixation of the 29113^CRISPRh^ allele. 20 heterozygous males and 20 heterozygous females for the 29113^CRISPRh^ allele were crossed with each other. The same cross was performed with 29113^hdrGFP/+^ individuals as control. **(A)** All adults, eggs and larvae were recorded. The resulting adults were used to set up the following generation. This procedure was continued until generation 7 (G7). The number of non-homing 29113^+/-^ adults was recorded as comparison until G2, once the population had exceeded the carrying capacity of the experimental set up. **(B)** Indels and substitutions seen at the predicted cleavage site for 29113^CRISPRh^ were assessed in 220 adults (G7) after repeated crossing of 29113^CRISPRh/+^ individuals for 7 generations. The gRNA binding site is underlined, with the PAM highlighted in bold. The Base highlighted in red shows a substitution that was also identified in the resistance assay.

## Discussion

Any strategy, from antibiotics to insecticides, that aims to impose a fitness load on a population inevitably leads to strong selection for resistance. For gene drives, the easiest pathway for resistance to occur is for an allele to arise that codes for a functioning gene that lacks the endonuclease recognition site, thereby blocking the biased inheritance of the gene drive element. These alleles may pre-exist in the population, arise through spontaneous point mutation, or arise through error-prone (non-HDR) DNA repair pathways that introduce small mutations in the repair of the double stranded break generated by the gene drive.

We developed a CRISPR homing gene drive targeting an ultra-conserved site in the viability gene *AGAP0029113* and showed that this gene is essential for development of both male and female mosquitoes. However, the invasive potential of the gene drive construct was hindered by high fitness costs in the heterozygous 29113^CRISPRh^/+ individuals compared to the wild-type strain. Fitness costs in gene drive carriers are not unusual and have been reported for other gene drive strains (1,19,20). A fitness cost, per se, in carriers will not prevent a gene drive increasing in frequency provided that the magnitude of its biased inheritance outweighs the fitness cost suffered in the carrier. However, in the case here, the fitness costs were severe enough to prevent the gene drive invading at all. One explanation for the fitness costs may be nuclease expression in the soma that converts the wild type allele to a null, thereby creating mosaic individuals with cells containing no functional copy of this essential gene. Mosaicism can derive from ‘leaky’ expression from the otherwise germline-restricted promoter or from parental deposition into the embryo. The germline promoter (from the gene *zpg*) used in this gene drive construct was shown previously to have minimal propensity to show these features, compared to alternative promoters, however we cannot exclude that position effects at the AGAP029113 locus may affect promoter specificity (17). An alternative explanation is that the target gene has a germline function in addition to its essential function in the soma, with the result that gene drive conversion to homozygosity in the germline leads to reduced fecundity. In our case it is difficult to disentangle such effects from general fitness effects since the competing hypothesis of somatic mosaicism could similarly lead to reduced fecundity.

Potentially such effects are more problematic for essential genes that are required during development in both sexes, compared to female fertility genes, since in the latter case heterozygous carrier males are unaffected. This is important for the phenotypic assessment of the suitability of future gene drive candidate genes required for viability in males and females and more emphasis should be placed on the restriction of gene drive activity to the germline. To look at the potential for gene drive resistance to arise at our chosen ultra-conserved target site, we devised both a single generation resistance assay and a population experiment designed to mimic the highest selection pressure. In both scenarios we recovered the same single point mutation at the target site. Possibly, the resistant allele we isolated might carry a small fitness cost which would be selected against in natural mosquito populations, which is why it was not detected within variation data from the 2784 *An. gambiae, An. coluzzii, Anopheles arabiensis* and hybrid individuals, collected from 19 countries in Africa as part of the Phase 3 MalariaGEN *Anopheles gambiae* 1000 Genomes Project (17). However, when faced with a gene drive, any small fitness cost is offset by the relative advantage the mutation confers against the gene drive allele. Therefore, even where genomic locations are ultra-conserved across various species and populations over a wide geographic area, their mutation may be tolerated under the selective pressure of gene drive. Further, different ultra-conserved sites could have very different functional constraints and selection pressure; studies have shown that even without any obvious functional constraints sequences can remain ultra-conserved between different species (21). Consequently, how to assess the potential of resistance development at any given conserved target site is of fundamental importance when considering gene drives of this type.

As a general rule, for a population suppression gene drive, the largest load is imparted on a population when the target is a gene essential for female fertility or viability, yet a considerable, though reduced, level of suppression can be achieved when the gene produces a recessive lethal phenotype that manifests in both sexes (22). Given that there may be a paucity of genes with functionally constrained target sites that are sufficiently robust to resistance it may be that the optimal target site choice will consider a balance between the suppressive load conferred by the gene drive and the intrinsic capability of its target site to tolerate resistant mutations.

Our mimicking of genotype fixation could be a useful experimental approach for swiftly assessing target site susceptibility, since it isolates resistant alleles faster than possible in population experiments of gene drive spread into wild-type populations - the selective advantage to a resistant allele over a wild type allele is highest when the likelihood of either allele finding themselves balanced against a gene drive allele is highest. A similar rationale applies in the resistance assay balancing the gene drive against a marked null allele (19,23) that we also employed here, however in this assay no measure is given of how strongly such potentially resistant alleles are selected or, indeed, of whether they are resistant to the gene drive nuclease. One feature of targeting a gene essential in early development, as we did here, is that it allows a much higher throughput of screening, due to the fact that selection happens during larval rearing, where rearing capacity is much less limited, meaning that all surviving adults have been through the selection bottleneck. In contrast, for a gene drive that targets female fertility in the adult, the vast majority of the adults will be homozygous for the gene drive and selection of a resistant allele only occurs in the small subset of females that have the opportunity to mate and contain at least one non-drive allele.

It will be essential to build on successes to date that have shown gene drives that can suppress populations robustly, without obvious selection for resistance (1). At scale, in a field setting and considering the vast population sizes of mosquitoes it is certain that such drives will need to be augmented by targeting multiple sites in the same gene, in order to greatly reduce the likelihood of multi-resistant alleles arising on the same haplotype (24–27). A similar effect can be achieved by releasing multiple gene drives that independently target separate loci (22,28). Moreover, using a similar logic, it could be prudent to have gene drives targeting genes in a range of different, independent biological pathways. In all cases, it will be essential to prioritise target site choice according not just to its suitability in terms of desired phenotypic effect when targeted, but its resilience to the generation of resistant alleles. Our results here show a pathway for how testing of ultra-conserved sites might proceed in order to evaluate these determinants for future gene drive designs in vivo.

## Materials and Methods

### Ethics statement

All animal work was conducted according to UK Home Office Regulations and approved under Home Office License PPL 70/6453.

### Generation of CRISPR and donor constructs

A CRISPR construct (p16510) containing a human-codon-optimized *Cas9* coding sequence (*hCas9*) under the control of the *zpg* promoter and U6::gRNA spacer cloning cassette was utilized as described previously and modified using Golden Gate cloning to generate a PolIII transcription unit containing the AGAP0*29113*-specific gRNA (1,2). The donor plasmid was assembled by MultiSite Gateway cloning (Invitrogen) and contained a *GFP* transcription unit under the control of the *3xP3* promoter enclosed within two reversible ϕC31 *attP* recombination sequences flanked both 5′ and 3′ by 2 kb sequence amplified using primer 29113 ex5 B1 f (<u>CAACCAAGTAGTTACTGTGCTC</u>) and 29113 ex5 B4 r (<u>GTCTTTTGTTGTTGTTCACGT</u>) and primer 29113 ex5 B3 f (<u>TGTAGGCCGTGATCGTGC</u>) and 29113 ex5 B2 r (<u>GCGACACCATACTCCGATG</u>) of the exon 5 target site in *AGAP029113*. To generate the 29113^CRISPRh^ allele a gRNA spacer (<u>GAACAACAACAAAAGACTGTAGG</u>) bearing complete homology to the intended target sequence was inserted by Golden Gate cloning into a CRISPR construct (p17410) as described previously (1,2). This vector contains a human codon-optimized Cas9 gene (hCas9) under control of the zpg promoter and a U6::gRNA cassette with a BsaI cloning site, as well as a visual marker (3xP3::RFP). This constructs sequence is flanked by attB recombination sites, which will allow the recombinase-mediated in vivo cassette exchange for the homing allele.

### Generation of the 29113^hdrGFP^ docking line and 29113 ^CRISPRh^ line

In order to generate the docking line, we injected 253 eggs of the *Anopheles gambiae* G3 strain with the 29113gRNA-modified p16510 plasmid (300ng/μl) and donor plasmid (300ng/μl) and obtained 7 transients that led to 6 transgenic individuals. Site-specific Integration was confirmed by PCR using primers binding the docking construct, 5′GFP-R (<u>TGAACAGCTCCTCGCCCTTG</u>) and a primer binding the genome outside of the homology arms: DL_29113 Ex5_F2 (<u>TTCCACCTCTCGCTCGTAGT)</u> and sequencing of the flanking sites. For the recombinase-mediated cassette exchange reactions a mix containing the CRISPR plasmid (200 ng/μl) and 400 ng/μl vasa2::integrase helper plasmid (29) was injected into embryos of the 29113^+/-^docking lines. Progeny from the outcross of surviving G_0_ individuals to WT were screened for the presence of RFP and the absence of GFP that should be indicative of a successful cassette-exchange event.

### Assessment of homing rate of CRISPR construct

To assess the ability of this 29113^CRISPRh^ construct to home at super-Mendelian rates, individuals heterozygous (29113^CRISPRh^/+) for the construct were crossed with wild type individuals and the progeny scored for inheritance of the CRISPR construct (via the proxy of RFP) from individual lays (N=15 per cross).

### Molecular characterisation of homozygotes and heterozygotes for the 29113^hdrGFP^ allele

Males and females heterozygote for the 29113^hdrGFP^ allele were crossed with each other and 60 L1-L2 larvae were analysed by Multiplex PCR using primer 3’-GFP-F (<u>GCCCTGAGCAAAGACCCCAA</u>), 5’-GFP R (<u>TGAACAGCTCCTCGCCCTTG</u>) and the primer DL_29113 Ex5_F2 (<u>TTCCACCTCTCGCTCGTAGT)</u>flanking the transgenic construct. The same PCR was repeated for 60 adult mosquitoes from the same cross.

### Fertility and fecundity data

Groups of a minimum of 40 virgin heterozygotes carrying the 29113^hdrGFP/+^ or 29113^CRISPRh/+^ genotype respectively were mated to an equal number of virgin wildtype mosquitoes for 5 days and the number of eggs and offspring was counted from individuals lays (N≥12). Females that failed to lay eggs and did not contain sperm in their spermathecal were excluded from the analysis.

### Larval to adult eclosion rate

Groups of a minimum of 40 virgin heterozygotes carrying the 29113^hdrGFP/+^ or 29113^CRISPRh/+^ genotype respectively were mated to an equal number of virgin wildtype mosquitoes for 5 days. 100 transgenic L1 larvae were selected and reared to adulthood for each experiment (N≥3).

### Population cage experiments with 20% starting frequency

First instar mosquito larvae heterozygous for the 29113^CRISPRh^ allele were mixed within 12 h of eclosion with age-matched WT larvae at a ratio of 1:5 in rearing trays at a density of 200 per tray (in ∼1 litre rearing water). The mixed population was used to seed three cages (36cm^3^) with 400 adult mosquitoes each. For three generations, each cage was fed after allowing 5 days for mating, and an egg bowl placed in the cage 48 h after a blood meal to allow overnight oviposition. A random sample of 450 eggs was selected. After allowing full eclosion, offspring were scored under fluorescence microscopy for the presence or absence of the RFP-linked *CRISPR*^*h*^ allele, then reared together in the same trays and then used to populate the next generation.

### Population model

To model the results of the population cage experiment, we use a simple deterministic model of a single, randomly mating population with two life stages (juveniles and adults), two sexes and discrete non-overlapping generations, following the structure of Beaghton et al (18), without considering parental deposition. We initially consider 3 alleles: wild-type (W), transgenic (T) and cleavage resistant (R). We model insertion of the transgene into a recessive gene required for survival to adulthood and assume all cleavage resistant alleles are non-functional, therefore T/T, T/R and R/R adults are inviable. For the 29113^CRISPRh^ transgene, cleavage of the W allele occurs in W/T individuals, followed by either homology directed repair, converting W to T, or non-homologous end joining, converting W to R. Cleavage and end joining rates were calculated from the proportion of progeny inheriting the 29113^*CRISPRh*^ allele (*d*) (F) = 0.927, (M) = 0.913 (Fig 2B) and the probability that non-homed alleles are R (*u*) = 0.266 (based on % of exposed but unmodified alleles from Fig 4B) where *e* = 2*d* − 1, cleavage = *e* + (1 − *e*)*u*, (F) = 0.893, (M) = 0.872 and end joining 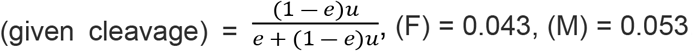. To model functional resistance, we extend the model to include a fourth allele (r), which is both cleavage resistant and fully functional. Here we allow a proportion (*p*) of non-homologous end joining products to be functional (r) and 1− *p* to be non-functional (R). To incorporate the heterozygote fitness costs identified from the phenotypic assays we consider female heterozygotes carrying the transgene to lay fewer viable eggs compared to wildtype: fecundity cost (1 - relative hatching rate) (F) = 0.216 (Fig 2D), and male and female heterozygotes to have reduced survival to adulthood: somatic cost (1 – relative eclosion rate) (M) = 0.378, (F) = 0.462, (Fig 2E). We also consider the case where heterozygotes carrying the transgene have additional somatic costs of other unknown factors: somatic cost (M) = 0.9, (F) =0.9. In both cases W/R and W/r individuals are assumed to be fully fit. When modelling the 29113^+/-^ line we assumed mendelian inheritance of the transgene and no heterozygous fitness costs. Following the experimental protocols, the frequency of transgenics was calculated at the juvenile stage, except for generation G^0^ which reflects that of the starting adult population.

### Small population cage experiment mimicking fixation of the drive

20 male and 20 virgin female adults heterozygous for the 23113^*CRISPRh*^ allele were placed in a small cage (25cm^3^). Females were blood-fed after 5 days and an egg bowl was placed in the cage 48h post bloodmeal. After allowing full eclosion the number of eggs and RFP+ and RFP-larvae was recorded. All offspring was reared to adulthood and crossed with each other. This was continued for 7 generations. The same procedure was repeated for adults heterozygous for the 29113^+/-^ allele for 2 generations. Offspring was screened for GFP+.

### Resistance Assay

400 males heterozygous for the 29113^CRISPRh^ allele (RFP+) were crossed to 400 females heterozygous for the 29113^+/-^ allele. To collect L1 for DNA extractions, 7000 individuals were screened for GFP using COPAS. Larvae that were GFP+ only or RFP/GFP+ (RFP+ only and larvae with no fluorescent marker were discarded) were pooled and frozen in ethanol at −20°C. To collect adults for DNA extraction large number of larvae were needed (as the majority would die before reaching adulthood). 10,000 L1 were screened for GFP+, and the sorted larvae were split into 20 trays of 500 individuals. A single tray was then reassessed for frequency of GFP+ and RFP/GFP+. This tray was then rescreened at L4 stage for frequency of GFP+, RFP/GFP+ and the rate of larval mortality. Surviving larvae were collected from all 20 trays and screened manually for GFP+. These were allowed to emerge as adults before collection and storage at −20°C for later DNA extraction.

### PCR of target site and deep sequencing analysis preparation

Pooled DNA extractions (minimum 90 adult mosquitoes or 3500 L1 larvae) were performed using the Wizard Genomic DNA Purification kit (Promega). A 349 bp locus containing the predicted on-target cleavage site was amplified with primers containing Illumina Nextera Transposase adapters (underlined), 29113-F (<u>TCGTCGGCAGCGTCAGATGTGTATAAGAGACAG</u>CTGCTTGCGGGAACACTGTTAG) and 29113-R (<u>GTCTCGTGGGCTCGGAGATGTGTATAAGAGACAG</u>TGTTAAGGAAGTCCCGACAGCG). PCR reactions with KAPA HiFi HotStart ReadyMix PCR Kit (Kapa Biosystems) were setup with 80ng genomic DNA. These PCR reactions, library preparation and deep sequencing were performed as previously described (1).

### Deep sequencing analysis

CRISPResso software v2.0.29 (30) was utilised for analysis of amplicon sequencing around the on-target site, with the parameter –q 30. Reads containing indels and substitutions 1bp either side of the predicted cleavage site were tallied as modified. Frameshift analysis was performed within CRISPResso2 using parameter –c to input the coding sequence.

### Statistical analysis

To compare the number of eggs laid by different strains, a quasi-Poisson generalised linear model (GLM) was fitted to the egg counts, with stains (5 levels, or categories) being the main effect. Quasi-binomial was used to explore the difference in the hatching rates across. The quasi family was adopted as the data exhibits overdispersion. A mixed-effect binomial GLM was run to study the eclosion rates for the various strains and across generations. A random effect was included for experiments with replicates.

## Acknowledgements

We are grateful for the helpful feedback on the manuscript by John Connolly and John Mumford. The authors are members of the Target Malaria not-for-profit research consortium, which receives core funding from the Bill & Melinda Gates Foundation (Grant INV006610 Target Malaria Phase II) and from the Open Philanthropy Project Fund (Grant O-77157).

## Supporting information

**Fig S1.**
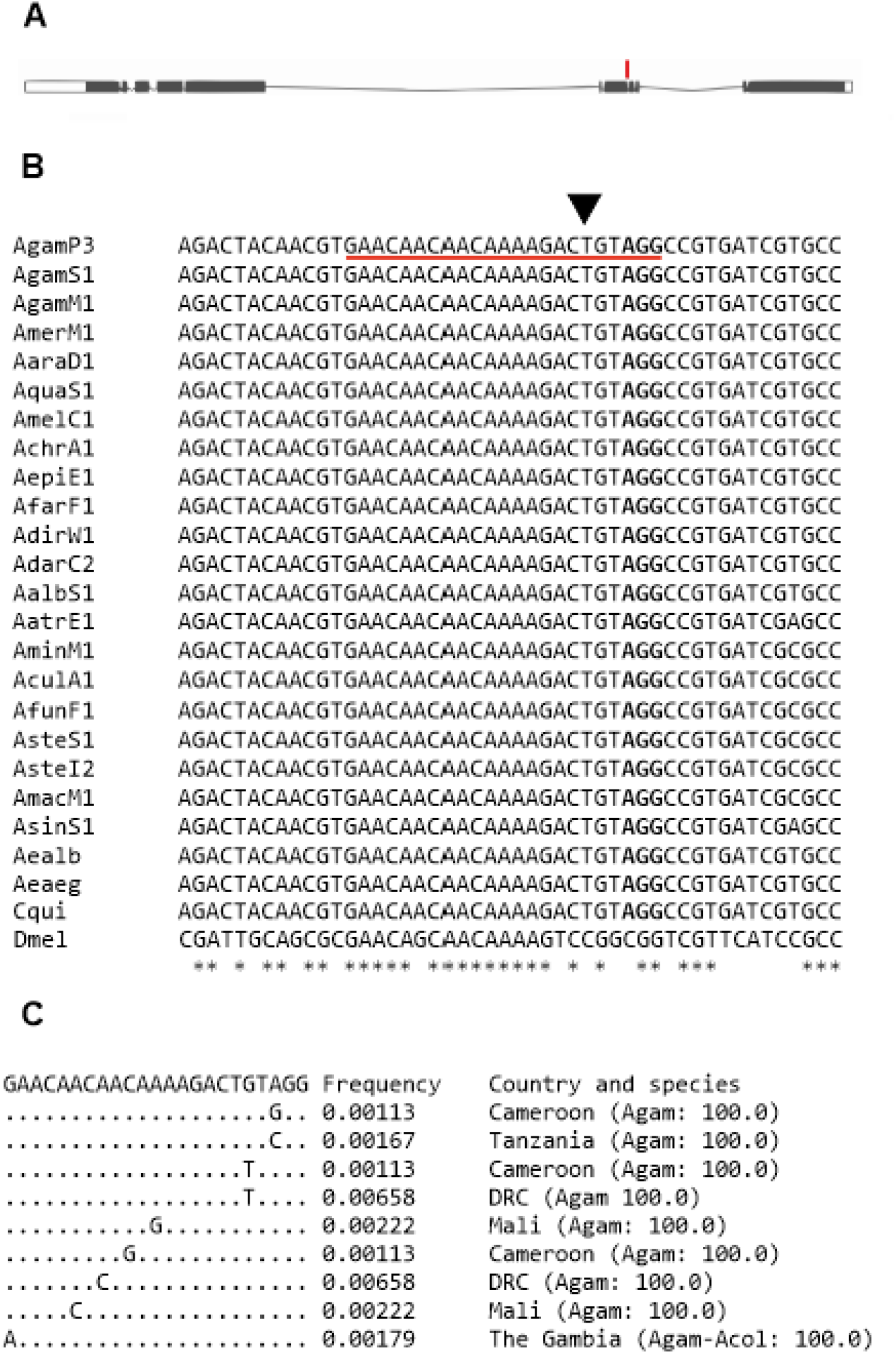
Target site selection in *AGAP029113*. **(A)** Schematic representation of the 12 exons of AGAP029113 and location of the target site in exon 5 (red bar). **(B)** Nucleotide alignment of exon 4 and 5 sections of AGAP029113 (Chromosome 2L: 2921441-2921488) belonging to *Anopheles gambiae/coluzzii* (AgamP3, AgamS1, AgamM1), *An. merus* (AmerM1), *An. arabiensis* (AaraD1), *Anopheles quadriannulatus* (AquaS1), *An. melas* (AmelC1), *Anopheles christyi* (AchrA1), *Anophles epiroticus* (AepiE1), *Anopheles minimus* (AminM1), *Anopheles culicifacies* (AculA1), *Anopheles funestus* (AfunF1), *Anopheles stephensi* (AsteS1, AsteI2), *Anopheles maculatus* (AmacM1), *Anopheles farauti* (AfarF1), *Anopheles dirus* (AdirW1), *Anopheles sinensis* (AsinS1), *Anopheles atroparvus* (AatrE1), *Anopheles darlingi* (AdarC2), *Anopheles albimanus* (AalbS1), *An. funestus* (AfunF1), *Ae. Albopictus* (Aealb), *Aedes aegypti* (Aeaeg), *Culex quinquefasciatus* (Cqui) and *Drosophila melanogaster* (Dmel) CRISPR gRNA target is underlined in red and the CRISPR-Cas9 cut site is indicated by a black arrowhead. The PAM sequence is indicated in bold letters. **(C)** Ag1000g phase 3 SNP data and frequency at the AGAP029113 target site sampled from 2784 wild *Anopheles* mosquitoes across Africa and 297 parents and progeny from 15 lab crosses. *Anopheles gambiae* (Agam), *Anopheles coluzzii* (Acol).

**Fig S2.**
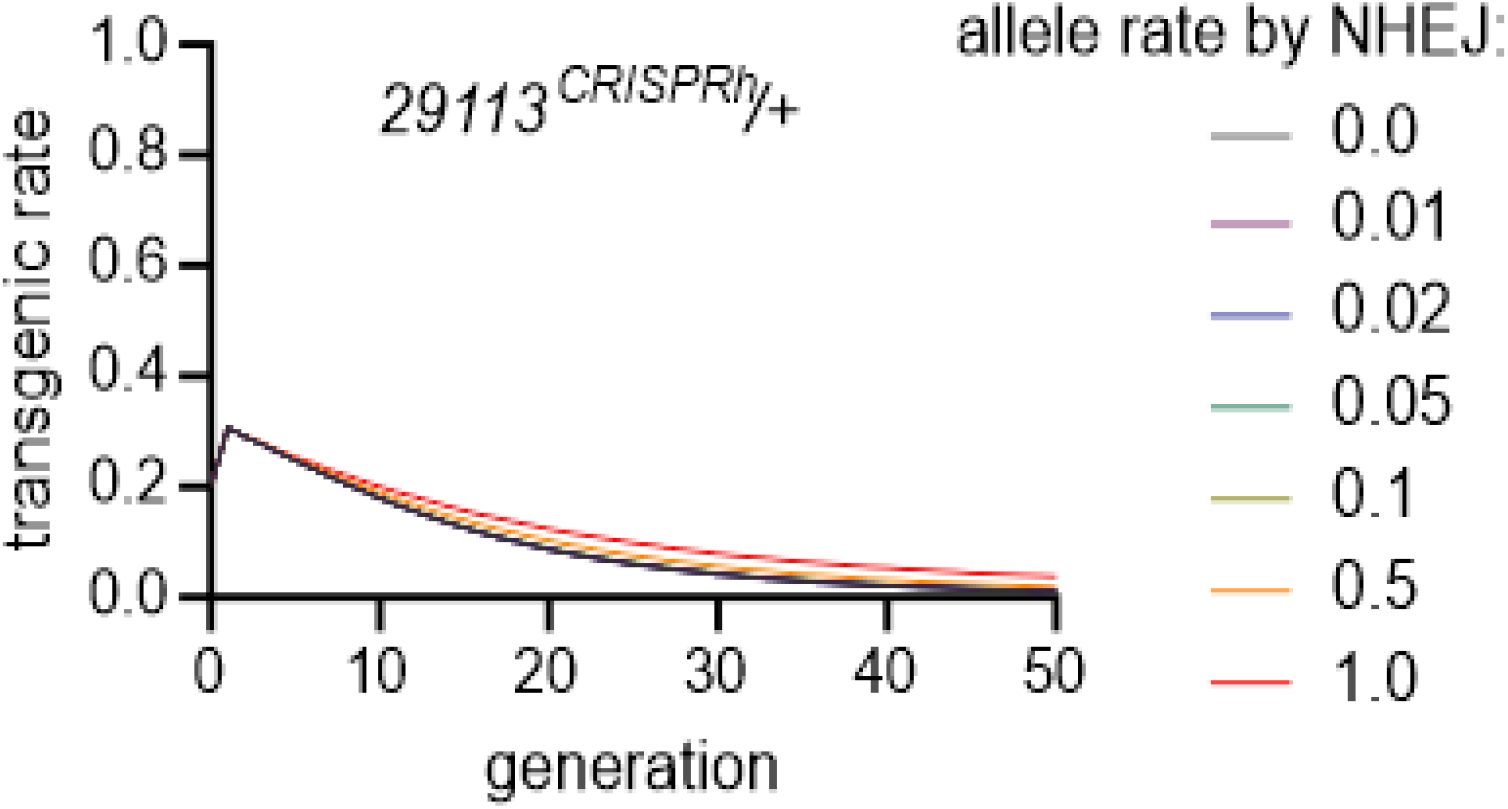
Deterministic prediction of the proportion of 29113^CRISPRh/+^ transgenics under observed fitness costs and varying probability of generation of a functional resistance (r) allele by NHEJ in the germline.

**Fig S3.**
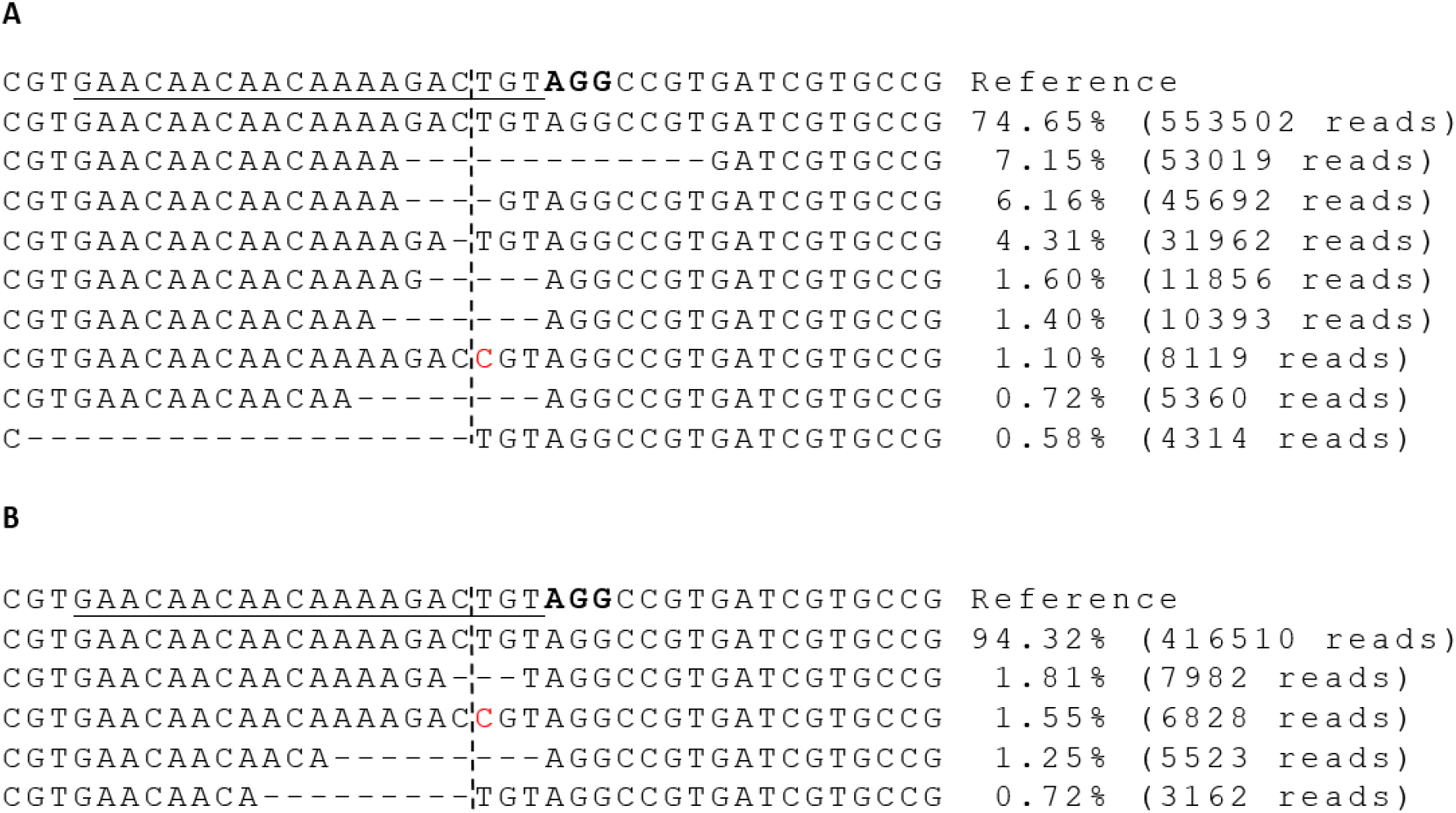
Indels and substitutions seen at the predicted cleavage site for 29113^CRISPRh^ in the resistance assay in larvae and adults. Crossing of heterozygous 29113^CRISPRh/+^ individuals to 29113^hdrGFP/+^ individuals, and subsequent screening of the offspring for GFP+, provided mosquitoes with a deficient 29113 allele, with the other allele being wild-type or containing indels generated by NHEJ repair. **(A)** L1 larvae (7000). **(B)** Adults (90). The gRNA binding site is underlined, with the PAM highlighted in bold. Horizontal dashes represent deletions whilst vertical dashes show the predicted cleavage site. Bases highlighted in red show substitutions, while bases in a red box represent insertions.

